# Coronin-1 controls directional cell rearrangement in *Drosophila* wing epithelium

**DOI:** 10.1101/2023.07.23.550233

**Authors:** Keisuke Ikawa, Souta Hiro, Shu Kondo, Shizue Ohsawa, Kaoru Sugimura

## Abstract

Directional cell rearrangement is a critical process underlying correct tissue deformation during morphogenesis. Although the involvement of F-actin regulation in cell rearrangement has been established, the role and regulation of actin binding proteins (ABPs) in this process are not well understood. In this study, we investigated the function of Coronin-1, a WD-repeat actin-binding protein, in controlling directional cell rearrangement in the *Drosophila* pupal wing. Transgenic flies expressing Coronin-1-EGFP were generated using CRISPR-Cas9. We observed that Coronin-1 localizes at the reconnecting junction during cell rearrangement, which is dependent on actin interacting protein 1 (AIP1), an actin disassembler and a known regulator of wing cell rearrangement. Loss of Coronin-1 function reduces cell rearrangement directionality and hexagonal cell fraction. These results suggest that Coronin-1 promotes directional cell rearrangement via its interaction with AIP1, highlighting the role of ABPs in the complex process of morphogenesis.

## Introduction

Cell rearrangement plays a critical role in morphogenesis (Takeichi, 2014). It involves shrinkage, reconnection, and elongation of the adherence junction (AJ) to alter the relative cell positions. The reorganization of the F-actin network by actin binding proteins (ABPs) is essential for remodeling the junction, as the AJ complex is tightly linked to the actomyosin network (Yap *et al*., 2015; Clarke and Martin, 2021). However, the precise molecular mechanisms governing F-actin dynamics during cell rearrangement remain unclear.

*Drosophila* pupal wing is an ideal model system to study the mechanisms of cell rearrangement. In the wing tissue, the direction of cell rearrangement is precisely regulated. Approximately 21–22 hours after puparium formation (h APF), the anterior-posterior (AP) junction shrinks, while the newly generated junction elongates along the proximal-distal (PD) axis, resulting in contraction-elongation of the wing tissue (Aigouy *et al*., 2010; Sugimura and Ishihara, 2013). Myosin-II (myo-II) and its regulatory molecules trigger junction shrinkage and elongation in the wing (Bardet *et al*., 2013). In addition, actin interacting protein 1 (AIP1) and cofilin, which disassemble F-actin, guide cell rearrangement by promoting actin turnover along the shrinking AP junctions (Ikawa and Sugimura, 2018). However, the role of other ABPs in cell rearrangement remains largely unexplored.

Cofilin relies on co-factors, including AIP1, for efficient actin filament severing (McCall *et al*., 2019; Nadkarni and Brieher, 2014). Coronin-1 (*Drosophila* gene name, *coronin*) is another cofactor of cofilin. It exerts bidirectional control over F-actin dynamics—it facilitates both cofilin-mediated F-actin severing and Arp2/3-mediated F-actin branching (Brieher *et al*., 2006; Gandhi *et al*., 2009). Notably, it has been reported that *coronin* mutants exhibit malformed wing shapes (Bharathi *et al*., 2004), suggesting cell rearrangement defects.

Here, we show that Coronin-1 controls directional cell rearrangement in the *Drosophila* pupal wing. Using CRISPR-Cas9 techniques, we generated transgenic flies expressing Coronin-1-EGFP to observe its localization and dynamics. The observations indicated that Coronin-1 is localized at the reconnecting junction during cell rearrangement and that this localization is dependent on AIP1. Moreover, the loss of Coronin-1 functions resulted in directional cell rearrangement defects. These findings suggest that AIP1 and Coronin-1 interplay governs directional cell rearrangement.

## Methods

### Generation of transgenic flies

The Coronin-1-EGFP knock-in line was generated by targeted integration of an EGFP sequence into the 3′ end of the *coronin* (*Drosophila coronin-1* gene) gene using CRISPR/Cas9. An EGFP knock-in cassette vector, pPExRF3, was constructed by replacing the Venus sequence with the EGFP sequence in pPVxRF3 (Kondo *et al*., 2020). The left and right homology arms of approximately 2000-bp were PCR-amplified from the genomic DNA of the standard *y*^*1*^; *cn*^*1*^ *bw*^*1*^ *sp*^*1*^ strain. Their sequences were designed such that EGFP was translated as an in-frame C-terminal fusion with the target protein. The reporter cassette excised from pPExRF3 and the left and right homology arms were assembled and cloned into a linearized pUC19 in a single enzymatic reaction using the In-Fusion Cloning Kit (Clontech Laboratories Inc., California, U.S.A). A gRNA vector was constructed by cloning a 20-bp target sequence surrounding the stop codon of *coronin* into pU6 (FlyCRISPR). A mixture of the donor vector (100 ng/μL) and the gRNA vector (100 ng/μL) were injected into fertilized eggs of *y*^*1*^, *w*^*1118*^; *attP40{nos-Cas9}/CyO*, which maternally express Cas9 protein (Kondo and Ueda, 2013); 100–200 eggs were injected, either in-house or by BestGene Inc. The surviving larvae were reared to adulthood and crossed to *y*^*1*^, *w*^*1118*^. Transformants in the F1 progeny were selected by eye-specific RFP expression from a *3xP3-RFP* marker gene in adults and a single transformant was used to establish a transgenic line. The *3xP3-RFP* marker was subsequently removed by first crossing them to *CyO-Crew*, a balancer chromosome carrying *hs-Cre* (Siegal and Hartl, 1996). The resulting males were then crossed to generic balancer lines, and their progeny were screened for loss of the eye-specific RFP expression. Leaky expression of Cre from the *hs-Cre* transgene in the absence of heat shock was sufficient to induce excision in almost all of the progeny.

### *Drosophila* genetics

The fly strains and a list of genotypes are summarized in the Supplementary Methods.

### Image analysis

#### Subcellular distribution of proteins

The quantification of signal intensities at rectangle-shaped myo-II cables (rsMC) was performed as described previously (Ikawa *et al*., 2023). Briefly, Z-projection was performed to extract the fluorescent signals on the AJ planes. The background signal of myo-II-mKate2 or Coronin-1-EGFP was subtracted using the ‘subtract background’ command (*r* = 50), and the signal intensity in manually selected regions of interest (ROIs) was measured using the ROI manager in ImageJ. The myo-II-mKate2 or Coronin-1-EGFP signal intensities at the rsMC were calculated using the ‘Plot Profile’ in ImageJ with a line width of 10 pixels.

#### Direction of cell rearrangement

Cell rearrangement was manually detected from time-lapse movies captured at 1-min intervals starting from 24 h APF at 25 °C (Ikawa and Sugimura, 2018).

#### Fraction of hexagonal cells

DE-cad-GFP images were skeletonized using ImageJ. The position and connectivity of vertices were extracted from the skeletonized images, and the polygonal distribution of cells was measured in OpenCV (Sugimura and Ishihara, 2013).

### Statistical analysis

P-values were calculated based on the Fisher’s exact test and Student’s t-test using EZR, a graphical user interface for R (Saitama Medical Center, Jichi Medical University, Saitama, Japan, https://www.jichi.ac.jp/saitama-sct/SaitamaHP.files/statmed.html) (Kanda, 2012).

## Results

### Coronin-1 is localized at the reconnecting junction

To investigate the dynamics of Coronin-1 during cell rearrangement, we used CRISPR-Cas9 techniques to generate a transgenic fly expressing Coronin-1-EGFP (Fig. 1B). Live imaging confirmed the expression of the EGFP signal in the transgenic flies (Fig. 1C). Furthermore, the EGFP signal was specifically reduced in the C-region of the wing upon the expression of dsRNA against *coronin* using the *ptc*-Gal4 driver (Fig. 1A, D). These results demonstrate the successful generation of a Coronin-1-EGFP fly.

**Figure 1.**
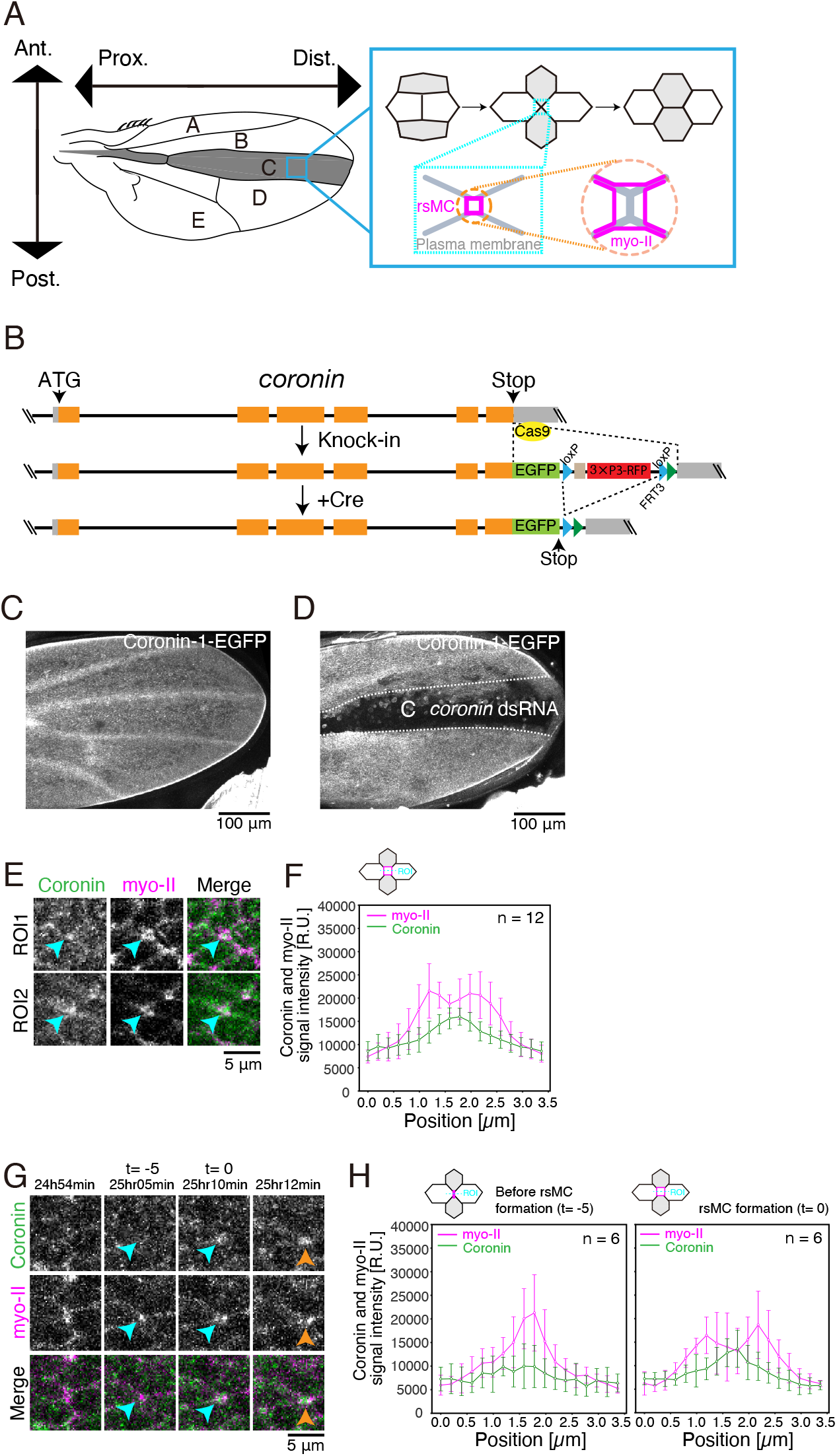
Endogenous Coronin-1 localizes at reconnecting junctions. (A) A schematic of the *Drosophila* wing and cell rearrangement during wing development. The vertical and horizontal directions are aligned with the anterior-posterior (AP) and the proximal-distal (PD) axes, respectively. The axes are conserved throughout all figures. *ptc*-Gal4 is expressed in the C region, shaded grey. Approximately 21–22 h after puparium formation (APF) and afterwards, wing cells shorten AP junctions and intercalate along the PD axis (Aigouy *et al*., 2010; Sugimura and Ishihara, 2013). During cell rearrangement, myo-II cables (magenta) are detached from short reconnecting junctions (designated as rectangle-shaped myo-II cables (rsMC); Ikawa *et al*., 2023). (B) Generation of an EGFP knock-in allele at the *coronin* locus. See Methods for details. (C, D) (C,D) Low magnification images of Coronin-1-EGFP of the *WT* and *coronin* RNAi wings at 24 h APF. (E)Images of Coronin-1-EGFP (gray in left panels and green in right panels) and myo-II-mKate2 (gray in middle panels and magenta in right panels). Arrowheads point to short, reconnecting junctions. (F)Quantifications of Coronin-1-EGFP and myo-II mKate2 signal intensities around the rsMC based on images in (E). (G) Time-lapse images of Coronin-1-EGFP (gray in upper panels, green in bottom panels) and myo-II-mKate2 (gray in bottom panels, magenta in bottom panels) during cell rearrangement. Blue and orange arrowheads indicate the AP and PD junctions, respectively. (H) Quantifications of Coronin-1-EGFP and myo-II mKate2 signal intensities around the rsMC based on timelapse images in (G). The number of regions of interest (ROIs) is indicated (F and H). The data are presented as mean ± SD (F and H). Scale bars: 100 μm (C and D) and 5 μm (E and G).

Using the Coronin-1-EGFP fly, we analyzed the subcellular localization of Coronin-1 during cell rearrangement. We observed a strong signal of Coronin-1-EGFP at short junctions (Fig. 1E). Previous studies have shown that during junction reconnection, the myo-II cables detach from the AJ and form rsMCs (magenta square in Fig. 1A; Ikawa *et al*., 2023). Our image analysis revealed a single peak of Coronin-1-EGFP between two peaks of myo-II-mKate2, indicating Coronin-1 accumulation within the rsMC (Fig. 1E, F). Furthermore, time-lapse analysis showed that the level of Coronin-1 increased at reconnecting junctions (Fig. 1G and H).

### *coronin* RNAi resulted in defects in directional cell rearrangement

We measured the angle of newly generated junctions following cell rearrangement to determine whether Coronin-1 is required for directional cell rearrangement in the wing (Fig. 2A). In *WT* wings, cell rearrangement was biased towards the PD axis, with ∼70% of the cell rearrangement occurring along this axis (Fig. 2B). In contrast, the proportion of PD cell rearrangement was reduced to ∼50% by RNAi of *coronin* (Fig. 2B). We next analyzed hexagonal cell packing, which is facilitated by directional cell rearrangement in the wing (Aigouy *et al*., 2010; Sugimura and Ishihara, 2013). The fraction of hexagonal cells decreased in *coronin* RNAi wings (75.6 ± 3.1% in *WT* and 57.8 ± 3.3% in *coronin* RNAi wings; Fig. 2C–E), consistent with the decrease in directional cell rearrangement. Collectively, these data indicate that Coronin-1 controls directional cell rearrangement, thereby supporting hexagonal cell packing in the wing.

**Figure 2.**
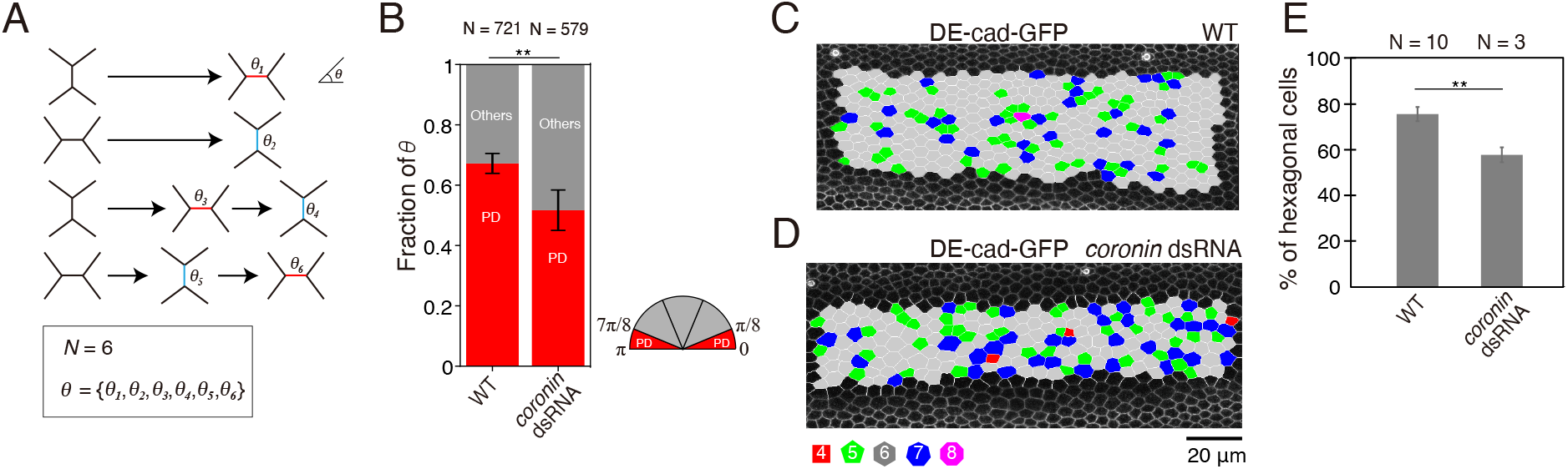
*coronin* RNAi results in defects in directional cell rearrangement. (A) Schematic of cell rearrangement analysis. We tracked individual junctions that appeared in a movie and measured their angle relative to the PD axes (*θ*) of newly generated junctions following cell rearrangement. (B) Quantification of the direction of cell rearrangement for each genotype based on time-lapse data captured at 24–27 h after puparium formation (APF) at 25°C (*WT, coronin* RNAi). The classification of *θ* is illustrated with a semicircle (red: PD, gray: others). *WT* timelapse data acquired by Ikawa and Sugimura (2018). (C, D) Images of DE-cad-GFP with the indicated genotypes (C: *WT* at 32 h APF, D: *coronin* RNAi at 32 h APF). Cells are colored according to the number of junctions (red, square; green, pentagon; gray, hexagon; blue, heptagon; and magenta, octagon). (E) The percentage of hexagonal cells at 32 h APF in the *WT* and *coronin* RNAi wings. WT data aquired by Ikawa and Sugimura (2018). The number of cell rearrangements (B) and wings (E) is indicated. The data are presented as mean ± SD (B and E). Fisher’s exact test: *WT* vs. *coronin* dsRNA, ** P < 0.001 (B) and Student’s t-test: *WT* vs. *coronin* dsRNA, ** P < 0.01 (E). Scale bars: 20 μm (C, D).

### AIP1 is required for the junctional localization of Coronin-1

Finally, we examined the dependence of Coronin-1 and AIP1 on their subcellular localization. It has been shown that AIP1 localizes at shrinking AP junctions and inside rsMCs (Fig. 3A; Ikawa and Sugimura, 2018; Ikawa *et al*., 2023). We found that AIP1 localization was largely unaffected in *coronin* RNAi wings. AIP1-GFP localized at the AP junctions and rsMCs with a moderate increase in the GFP signal intensity (Fig. 3B). In contrast, RNAi of *flare* (*flr*; *Drosophila aip1* gene) severely disrupted the junctional localization of Coronin-1 and induced cytoplasmic patch formation (Fig. 3C and D). These results suggest differential requirements for cofilin cofactors according to their subcellular localization: whereas AIP1 can localize along the remodeling AP junctions in the absence of *coronin* function, AIP1 supports the junctional localization of Coronin-1.

**Figure 3.**
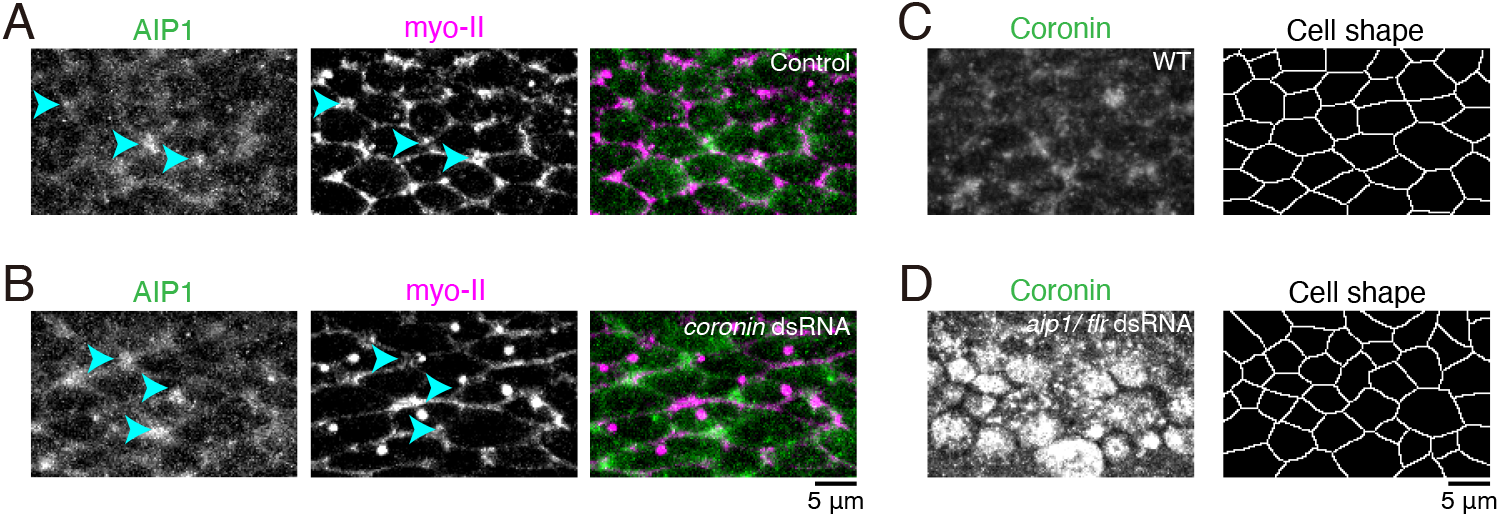
Coronin-1 localization is dependent on AIP1. (A, B) Images of AIP1-GFP (gray in left panels and green in right panels) and myo-II-mKate2 (gray in middle panels and magenta in right panels) in control (A) and *coronin* RNAi (B) wings at 24 h APF. (C, D) Images of Coronin-1-EGFP in *WT* (C) and *aip1*/*flr* RNAi (D) wings at 24 h APF. Cell shape is depicted in the right panels. Scale bars: 5 μm (A–D).

## Discussion

During cell rearrangement, F-actin, which is tightly linked to the junction structure, undergoes reorganization alongside junction remodeling (Yap *et al*., 2015; Clarke and Martin, 2021). In addition, F-actin remodeling has been implicated in the dissipation of excess stress along the shrinking junctions (Charras and Yap, 2018). ABPs play a critical role in controlling F-actin remodeling in eukaryotic cells (Munjal and Lecuit, 2014). However, the specific mechanisms and ABPs involved in regulating cell rearrangement remain unclear. In the present study, we demonstrated that Coronin-1, a cofilin cofactor, is localized at reconnecting junctions and governs directional cell rearrangement in the *Drosophila* wing. Moreover, we found that AIP1 promotes the junctional localization of Coronin-1. Exploring the interplay between cofilin, AIP1, and Coronin-1 is expected to provide insights into the mechanisms underlying F-actin remodeling during cell rearrangement.

Several *in vitro* studies have demonstrated that Coronin-1 promotes F-actin branching by Arp2/3 at the barbed ends, while both Coronin-1 and AIP1 promote F-actin severing by cofilin at the pointed ends (Brieher *et al*., 2006; Gandhi *et al*., 2009; Jansen *et al*., 2015). These studies have indicated that Coronin-1 promotes the localization of AIP1 and cofilin at the pointed ends (Gandhi *et al*., 2009; Jansen *et al*., 2015), which is inconsistent with our observation that the localization of AIP1 in the wing tissue was unaffected by *coronin* RNAi. There are two possible explanations for this inconsistency. First, *in vitro* experiments have shown that AIP1 and cofilin can still localize at the pointed ends, albeit to a lesser extent, in the absence of Coronin-1 (Ngo *et al*., 2015; Nadkarni and Brieher, 2014), suggesting that *coronin* RNAi may not significantly affect the subcellular localizations of AIP1 in wing tissue. Second, the function of Coronin-1 in F-actin branching may outweigh its role in F-actin severing in the wing tissue. Considering the significance of F-actin disassemblers in wing cell rearrangement (Ikawa and Sugimura, 2018), the former explanation appears more plausible. To address the inconsistencies between *in vitro* and *in vivo* studies, it is crucial to uncover the molecular mechanisms through which cofilin, AIP1, and Coronin-1 regulate F-actin remodeling during cell rearrangement.

Given that AIP1 and Coronin-1 are involved in other cellular processes, such as cell division and extrusion (Chen *et al*., 2015; Michael *et al*., 2016), studying the function of ABPs in these processes offers promising avenues for future research.

## Acknowledgments

The authors would like to thank Yohanns Bellaïche, Yang Hong, Roger Kares, and the Bloomington Stock Center; Miho Aruga, Kyoko Komano, and Risa Matsui for technical assistance; and the iCeMS Analysis Center for imaging equipment. This study was financially supported by a JSPS KAKENHI Grant (17K15125), the AMED PRIME program (20gm5810025h9904), the Sumitomo Foundation (200303), and the Takeda Science Foundation to K.S., a JSPS KAKENHI Grant (19K16139) and the Uehara Memorial Foundation (202110172) to K.I., a JSPS KAKENHI Grant (20H03246) to S.K., and a JSPS KAKENHI Grant (20H05945) to S.O.

## Author contributions

K.I. and K.S. designed the research. K.I. and S.H. performed the experiments. K.I., K.S and S.H. analyzed the data. S.K. generated transgenic flies. K.I. drafted the manuscript. K.I., K.S, S.O. and S.K revised the manuscript. All authors approved the final manuscript.

## Supplementary Methods

### Image collection

To prepare the *Drosophila* pupal wing samples for image collection, pupae at around 24 hour after puparium formation (h APF) were fixed to double-sided tape and the pupal case above the left wing was removed (Guirao *et al*., 2015, Ikawa and Sugimura, 2018). The pupae were then placed on a small drop of water or Immersol W 2010 (Zeiss 444969-0000-000) in a glass bottom dish with the left side facing downward. The fixed time-point and timelapse images of Coronin-1-EGFP were acquired using a confocal microscope (LSM900; Zeiss) equipped with a 63×/NA1.2 C-Apochromat water-immersion objective at room temperature. Images for the analysis of cell rearrangement were acquired using an inverted confocal spinning disk microscope (Olympus IX83 combined with Yokogawa CSU-W1) equipped with an iXon3 888 EMCCD camera (Andor), an Olympus 60×/NA1.2 SplanApo water-immersion objective and a temperature control chamber (TOKAI HIT), using IQ 2.9.1 (Andor) (Guirao *et al*., 2015). Images for the analysis of hexagonal cell packing were acquired using an inverted confocal microscope (A1R; Nikon) equipped with a 60×/NA1.2 Plan Apochromat water-immersion objective at 25°C. After imaging, we confirmed that the pupae survived to at least the pharate stage.

### *Drosophila* genetics

The flies used in the present study were *flr-GFP* (Flytrap #CA07499) (Buszczak *et al*., 2007), *DE-cad-GFP* (Huang *et al*., 2009), *sqhp-sqh-mKate2×3* (Pinheiro *et al*., 2017), *ptc-Gal4, UAS-flr dsRNA* (VDRC #v108422), *UAS-coro dsRNA* (BDSC #40841), *coro-EGFP* (this study). Fly genotypes and culture conditions are summarized below.

**Supplementary table 1.** Fly genotypes and culture conditions.

| Genotype | Culture condition | Figure |
| --- | --- | --- |
| *coro-EGFP/sqh-mKate2×3* | Crossed and observed at 25°C (24 h APF) | Fig. 1C, E, F, G and H |
| *ptc-Gal4, coro-EGFP/UAS-coro dsRNA* | Flies were crossed at 17°C. White pupae were piked up and observed at 25°C (24 h APF) | Fig. 1D |
| *DE-cad-GFP* | Flies were crossed at 17°C. white pupae were piked up and observed at 25°C (24 h APF) | Fig. 2B and C |
| *ptc-Gal4, DE-cad-GFP/DE-cad-GFP, UAS-coro dsRNA* | Flies were crossed at 17°C. white pupae were piked up and observed at 25°C (24 h APF) | Fig. 2B and D |
| *ptc-Gal4, sqh-mKate2×3/+; Flare-GFP/+* | Flies were crossed at 17°C. white pupae were piked up and observed at 25°C (24 h APF) | Fig. 3A |
| *ptc-Gal4, sqh-mKate2×3/UAS-coro dsRNA; Flare-GFP/+* | Flies were crossed at 17°C. white pupae were piked up and observed at 25°C (24 h APF) | Fig. 3B |
| *coro-EGFP/sqh-mKate2×3* | Crossed and observed at 25°C (24 h APF) | Fig. 3C |
| *ptc-Gal4, coro-EGFP/UAS-flr dsRNA* | White pupae were picked up and observed at 17°C. (58 h APF at 17°C) | Fig. 3D |

